# Noncanonical contributions of MutLγ to VDE-initiated crossovers during *Saccharomyces cerevisiae* meiosis

**DOI:** 10.1101/510297

**Authors:** Anura Shodhan, Darpan Medhi, Michael Lichten

## Abstract

In *Saccharomyces cerevisiae*, the meiosis-specific axis proteins Hop1 and Red1 are present nonuniformly across the genome. In a previous study, the meiosis-specific *VMA1-derived* endonuclease (VDE) was used to examine Spo11-independent recombination in a recombination reporter inserted in a Hop1/Red1-enriched region (*HIS4*) and in a Hop1/Red1-poor region (*URA3*). VDE-initiated crossovers at *HIS4* were mostly dependent on Mlh3, a component of the MutLγ meiotic recombination intermediate resolvase, while VDE-initiated crossovers at *URA3* were mostly Mlh3-independent. These differences were abolished in the absence of the chromosome axis remodeler Pch2, and crossovers at both loci become partly Mlh3-dependent. To test the generality of these observations, we examined inserts at six additional loci that differed in terms of Hop1/Red1 enrichment, chromosome size, and distance from centromeres and telomeres. All six loci behaved similarly to *URA3*: the vast majority of VDE-initiated crossovers were Mlh3-independent. This indicates that, counter to previous suggestions, meiotic chromosome axis protein enrichment is not a primary determinant of which recombination pathway gives rise to crossovers during VDE-initiated meiotic recombination. In *pch2*Δ mutants, the fraction of VDE-induced crossovers that were Mlh3-dependent increased to levels previously observed for Spo11-initiated crossovers in *pch2*Δ, indicating that Pch2-dependent processes play an important role in controlling the distribution of factors necessary for MutLγ-dependent crossovers.

## Introduction

During meiosis, the crossover products of recombination form stable links between homologous chromosomes of different parental origin (homologs), to enable their proper segregation during the meiotic divisions (reviewed by Zickler and Kleckner 1999; Whitby 2005). Meiotic recombination is initiated by DNA double strand breaks (DSBs) that are formed by the meiosis-specific Spo11 protein (Bergerat *et al.* 1997; Keeney 2001). In budding yeast, Spo11 DSBs are unevenly distributed in the genome. Most DSB-rich regions correlate with domains that are enriched for the meiosis-specific chromosome axis proteins, Red1 and Hop1, which play an important role in DSB formation (Hollingsworth and Ponte 1997; Blat *et al.* 2002; Pan *et al.* 2011; Panizza *et al.* 2011; Smagulova *et al.* 2011; Baker *et al.* 2014). The non-uniform distribution of Hop1 is maintained by Pch2, a hexameric AAA+ ATPase (Chen *et al.* 2014). In *pch2* mutants, Hop1 persists longer and is more uniformly distributed on chromosomes; this is accompanied by a delay in meiotic progression and changes in the distribution of late-forming DSBs and COs (Börner *et al.* 2008; Joshi *et al.* 2009; Zanders and Alani 2009; Lambing *et al.* 2015; Subramanian *et al.* 2016; Subramanian *et al.* 2018).

Meiotic DSBs are also important for homolog colocalization, pairing and synapsis (Keeney *et al.* 1997; Romanienko and Camerini-Otero 2000; Baudat *et al.* 2013). Current thinking is that most DSBs are repaired either by a synthesis-dependent strand annealing pathway that forms non-crossovers (NCOs), or by a pathway that forms double Holiday junction (dHJ) intermediates that are resolved as crossovers (COs) by the MutLγ (Mlh1-Mlh3 and Exo1) meiosis-specific resolvase (Schwacha and Kleckner 1994; Wang *et al.* 1999; Khazanehdari and Borts 2000; Kirkpatrick *et al.* 2000; Tsubouchi and Ogawa 2000; Allers and Lichten 2001b; Allers and Lichten 2001a; Hoffmann *et al.* 2003; Argueso *et al.* 2004; Bishop and Zickler 2004; Nishant *et al.* 2008; Zakharyevich *et al.* 2010; Al-Sweel *et al.* 2017). In budding yeast, COs and NCOs are formed at similar levels, suggesting that roughly equal fractions of DSBs are repaired by these two pathways (Martini *et al.* 2006; Mancera *et al.* 2008). Apart from these two major pathways, a minor pathway uses mitotic resolvases (structure-selective nucleases, SSNs: Mus81-Mms4, Yen1 and Slx1-4) to form both NCOs and COs (de los Santos *et al.* 2003; Argueso *et al.* 2004; Lynn *et al.* 2007; Jessop and Lichten 2008; De Muyt *et al.* 2012; Zakharyevich *et al.* 2012; Agostinho *et al.* 2013; Oke *et al.* 2014). while the proteins and enzymatic activities contributing to each of these pathways has been the subject of considerable study (reviewed by Ehmsen and Heyer 2008; Hunter 2015; Manhart and Alani 2016), the question of what roles local chromosome environment might play in pathway choice remains much less explored. (Medhi *et al.*2016) addressed this question using a meiosis-specific endonuclease, VDE, that cleaves a recognition sequence (VRS) at high efficiency regardless of chromosomal context (Gimble and Thorner 1992; Gimble and Thorner 1993; Nogami *et al.* 2002; Fukuda *et al.* 2003; Medhi *et al.* 2016; this work). Like Spo11 DSBs, VDE DSBs are processed to form single-stranded overhangs that recruit the Rad51 and Dmc1 proteins that perform strand invasion and homology search (Bishop *et al.* 1992; Fukuda *et al.* 2003; Fukuda and Ohya 2006). Medhi *et al.* (2016) inserted a VRS-containing recombination reporter at two loci: *HIS4*, present in a region with high levels of both Spo11 DSBs and Hop1 binding; and *URA3*, in a region with low levels of Spo11 DSBs and Hop1 binding (Pan *et al.* 2011; Panizza *et al.* 2011). Most COs at *HIS4* were Mlh3-dependent, while COs at *URA3* were Mlh3-independent. In *pch2*Δ mutants, Hop1 occupancy at *HIS4* was reduced, as were the fraction of COs that were Mlh3-dependent, while at *URA3* the fraction of COs that were Mlh3-dependent increased. Based on these findings, Medhi *et al.* suggested that the local chromosome structure, in particular levels of Hop1 enrichment, may be an important determinant of CO pathway choice.

To test the generality of the above suggestion, we inserted the same VRS recombination reporter at six new loci with varying Hop1 occupancy in their vicinity and found that VDE-initiated meiotic COs at all six new loci were predominantly Mlh3-independent. Moreover, as previously seen for inserts at *URA3* (Medhi *et al.* 2016), *pch2Δ* mutation increased the fraction of COs that were Mlh3-dependent. These results indicate that the local level of Hop1 enrichment is not the sole determinant of CO pathway choice in VDE-induced meiotic recombination. They also suggest that, at most loci, VDE DSBs are repaired differently than are Spo11 DSBs.

## Materials and Methods

### Yeast strains

All strains (Table S2) used in this study are of SK1 background (Kane and Roth 1974), and were constructed by transformation or genetic crosses. The recombination reporter cassette with the VRS (cleavable) or VRS-103 (uncleavable) site in the *ARG4* gene (Medhi *et al.* 2016) were inserted by ends-out transformation (for VRS-containing inserts and for VRS-103 inserts at *FIR1* and *HSP30*, Figure S1A) or by ends-in transformation (for VRS-103 constructs at *CCT6, RIM15, IMD3* and *TRK2*, Figure S1B) at six different locations (primers used are listed in Table S1). Ends-in transformation was used for inserts at divergently transcribed loci to minimize effects on expression caused by disruption of 5’ untranslated regions. Transformation was performed with overlapping DNA fragments as illustrated in Figure S1. The VRS-arg4 and VRS-103-arg4 constructs are 5.5kb and 8.6kb long, respectively, with ~3kb sequence homology around the VRS site. This size difference, along with HindIII site differences, enables the detection of the parental and recombinant chromosomes on Southern blots (see Figure 2, below).

### Growth and sporulation

Strains were grown in pre-sporulation SPS medium and transferred to sporulation medium as described (Goyon and Lichten 1993), with the inclusion of 10μM CuSO_4_ in sporulation medium to induce VDE expression (Medhi *et al.* 2016). DNA samples were collected and processed as described (Allers and Lichten 2000; Jessop *et al.* 2005; Jessop *et al.* 2006).

### DNA extraction and southern hybridization

DNA was extracted from samples using the CTAB extraction method (Allers and Lichten 2000; Oh *et al.* 2009). Genomic DNA was digested with *Hind*III or *Hind*III and PI-*Sce*I, run on agarose gels, blotted, probed and analyzed as described (Medhi *et al.* 2016).

### Cytology

Cells were collected, stained with DAPI, and scored by epifluoresence microscopy to follow nuclear divisions as described (Kaur *et al.* 2018).

### Statistical analysis

GraphPad Prism was used for comparisons of mean values, using t-tests with the Holm-Sidak correction for multiple comparisons.

### Data availability

Strains and plasmids are available upon request. The authors affirm that all data necessary for confirming the conclusions of this article are represented fully within the article, tables, figures, supplementary figures and supplementary tables. Underlying data for all graphs will be made available in Supplementary File 1 upon publication.

## Results and Discussion

### VDE-initiated COs are Mlh3-independent at most insert sites

To further test the hypothesis that Hop1-enrichment determines the MutLγ-dependence of meiotic CO formation, six new sites were selected for VRS reporter insertion, one *(HSP30)* with regional Hop1 levels similar to those at *URA3*, and five (CCT6, *FIR1, RIM15, TRK2* and *IMD3)* with regional Hop1 levels similar to those at *HIS4* (Figure 1). Since it has been previously shown that Spo11-DSBs are reduced near centromeres and telomeres (Pan *et al.* 2011) and CO formation is regulated differently on longer and shorter chromosomes (Joshi *et al.* 2009; Zanders and Alani 2009), the new sites were selected such that they were on chromosomes of different sizes and were at varying distances from centromeres and telomeres. At each site, recombination products can be differentiated on Southern blots (Figure 2A, B), as was previously used to quantify DSBs, COs and NCOs (Medhi *et al.* 2016).

**Figure 1.**
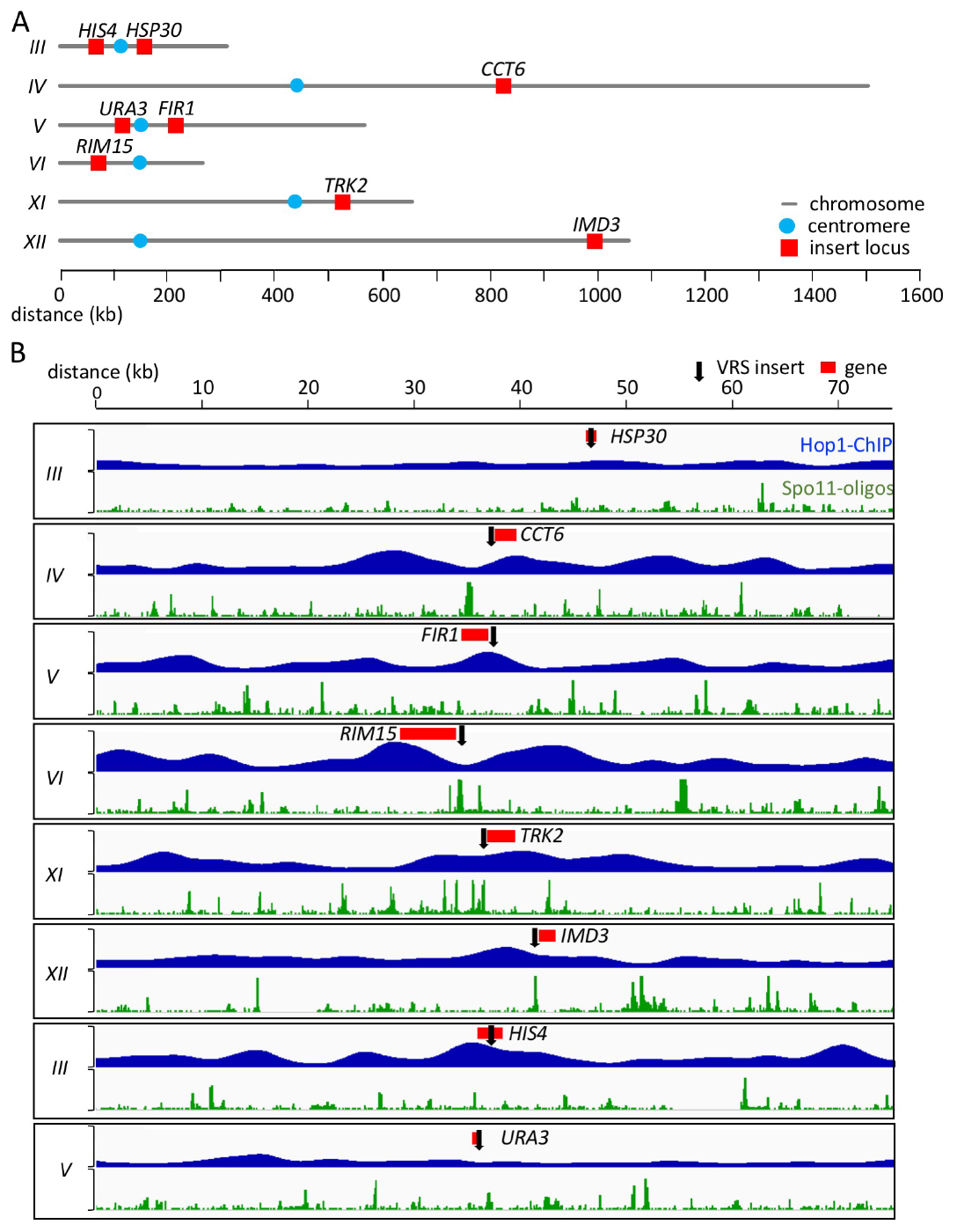
Insert loci examined. (A) Locations of insert loci are illustrated (red). Blue circles denote centromere locations. (B) Maps of regions surrounding insert loci. Red—coding region of gene used to identify each insert; black arrow—site of VRS insert. Blue plots show relative Hop1 occupancy levels in mid meiosis, using smoothed ChiP-chip data from (Pannizza *et al.* 2011); vertical scale = 0-7, decile-normalized ChiP/WCE. Green plots show relative DSB levels, using Spo11-oligo reads from Pan *et al.* (2011); vertical scale = 0-15 hits per million/base-pair.

**Figure 2.**
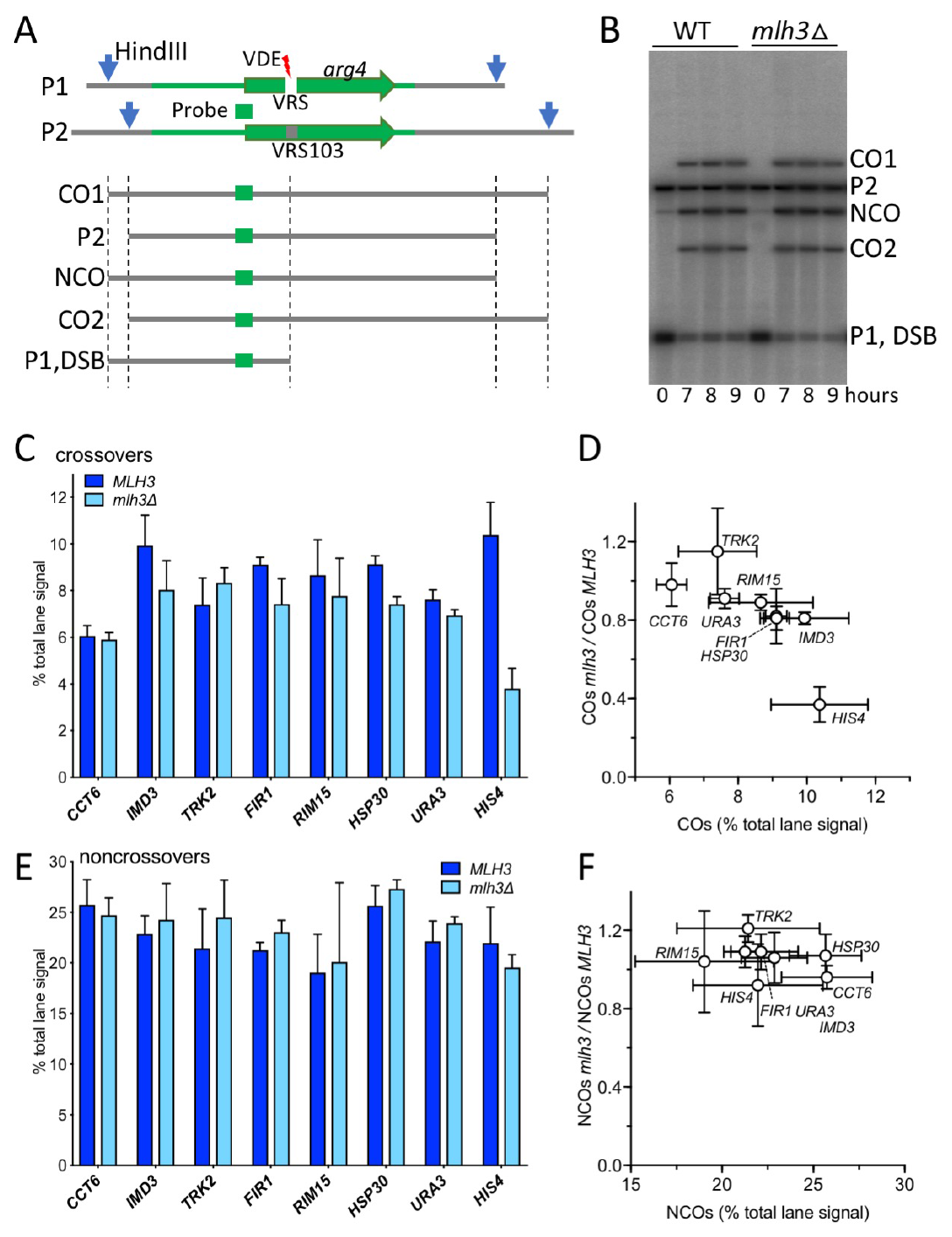
VDE-initiated crossovers at most loci are MutLγ-independent. (A) Strategy for detection of VDE-initiated COs and NCOs. A cartoon of the VRS and VRS-103 inserts is shown, illustrating : white box—VRS sequences; blue arrows*—Hind*III restriction sites; green lines—sequences shared between the two inserts, with *ARG4* coding sequences shown as a green arrow; green box—sequences used for Southern blot probes. Digestion with *Hind*III and PI-*Sce*I (VDE) distinguishes parental (P1 and P2), CO and NCO products. VDE-cut inserts are not distinguished from parent P1 on these digests, but can be distinguished in digests with HindIII alone (Medhi *et al.* 2016). (B) Representative Southern blot containing DNA from strains with inserts at *RJM15.* (C) VDE-initiated COs in *MLH3* and *mlh3*Δ cells. CO frequencies, average signal of CO1 and CO2 for 8 and 9 h samples from three independent experiments for inserts at *HIS4* and from two independent experiments for inserts at all other loci. Data for inserts at *URA3* and for two experiments with inserts at *HIS4* are from Medhi *et al.*, (2016). (D) fraction of COs that are MutLγ-independent (ratio of CO frequencies in *mlh3*Δ versus *MLH3)*, plotted as a function of CO frequencies in *MLH3* strains. CO frequencies in *MLH3* and *mlh3*Δ differ significantly only for inserts at *HSP30* and *HIS4* (adjusted *p* values of 0.003 and 0.0001, respectively) (E,F) VDE-initiated NCOs, details as in (B) and (C); frequencies in *MLH3* and *mlh3*Δ do not differ significantly at any locus. Error bars in all panels denote standard deviation.

Meiotic progression of all WT and *mlh3*Δ strains was similar, with most cells completing the first meiotic division by 7-8h post-induction (Figure S2A). In addition, VDE-initiated DSBs appeared and disappeared with levels and timing similar to those previously seen at *HIS4* and *URA3* (Figure S2B; Medhi *et al.* 2016).

COs in VRS inserts ranged from ~6% of total lane signal at *CCT6* to ~10.3% at *HIS4* (Figure 2C). As previously reported (Medhi *et al.*2016), NCOs were recovered in substantial excess over COs at all insert loci (Figure 2E), with NCO/CO ratios ranging from 2.1 to 4.8 (mean = 3.1 ± 0.8; Figure S2C). The marked excess of NCOs over COs seen for VDE-initiated events differs from what is seen with Spo11-initiated events, where COs and NCOs are produced at similar levels (Martini *et al.* 2006; Mancera *et al.* 2008; Zakharyevich *et al.* 2012). In contrast to what was seen for VRS inserts at *HIS4*, where COs were reduced dramatically in *mlh3*Δ mutants (to ~40% of wild-type levels), COs in the same sequences inserted at all other loci were only modestly affected, with COs in *mlh3*Δ ranging from ~80% to ~115% of wild type (mean = 91 ± 12%; Figure 2D); NCOs were similarly unaffected (Figure 2E, F). These results indicate that, in contrast to Spo11-initiated COs, which are reduced about 2-fold in *mlh3*Δ mutants (Wang *et al.* 1999; Khazanehdari and Borts 2000; Kirkpatrick *et al.* 2000; Tsubouchi and Ogawa 2000; Hoffmann *et al.* 2003; Argueso *et al.* 2004; Nishant *et al.* 2008; Al-Sweel *et al.* 2017; Chakraborty *et al.* 2017), most COs at the VDE break sites are formed independent of MutLγ, irrespective of the chromosome size, distance from centromere or telomere, or Hop1-enrichment in their vicinity. Thus, at most insert loci in otherwise wild-type cells, VDE-initiated recombination differs from Spoil-initiated recombination and more closely resembles mitotic recombination, in that NCOs are in excess over COs (Esposito 1978; Lichten and Haber 1989; Ira *et al.* 2003; Dayani *et al.* 2011) and, with the exception of those formed in inserts at *HIS4*, VDE-initiated COs are largely MutLγ-independent.

### *IncreasedMlh3-(lependence of VDE-initiated COs in* pch2Δ *mutants*

In *pch2* mutants, meiotic axis proteins are redistributed, with less pronounced differences in Hop1 occupancy distributions measured either cytologically (Börner *et al.* 2008; Joshi *et al.* 2009) or by chromatin-immunoprecipitation (Medhi *et al.* 2016). Previously, we found that the absence of Pch2 did not substantially alter overall NCO or CO levels at *HIS4* and *URA3*, but the Mlh3-dependence of CO formation was affected at both loci, with Mlh3-independent COs increasing at *HIS4* and decreasing at *URA3.* Because the six new VRS insert loci studied here are similar to *URA3*, in that most VDE-initiated COs are Mlh3-independent, we wanted to see if COs at these loci also displayed increased Mlh3-dependence in *pch2* Δ mutants.

Consistent with previous findings (Börner *et al.* 2008), meiotic divisions were delayed in *pch2 S.* and *pch2* Δ *mlh3* Δ mutants relative to wild type (Figure S2A). Frequencies of NCOs at all eight VRS insert loci in the *pch2* Δ were similar to those seen in wild type (Figure 3C; *pch2*Δ/*PCH2* = 111 ± 10%), as were COs (Figure 3A; *pch2 \ ľ(‘H2 =* 113 ± 16%). Loss of Mlh3 did not substantially affect NCOs in the absence of Pch2 (Figure 3C; *pch2* Δ *mlh3* Δ/*pch2*Δ *MLH3* = 114 ± 14%). However, in *pch2* Δ *mlh3* Δ double mutants, COs were reduced 20-35% relative to *pch2* Δ *MIH3* (Figure 3B; average *pch2* Δ *mlh3* Δ/*pch2*Δ = 74 ± 7%), as was previously observed for inserts at *URA3* and *HIS4* (Medhi *et al.* 2016). A quantitatively similar MutLγ-dependence has also been observed for Spoil-initiated COs in *pch2* Δ mutants, both genome-wide (*pch2* Δ *mlh3* Δ / *pch2* Δ = 73% (Chakraborty *et al.* 2017) and for individual genetic intervals (*pch2*Δ / *pch2* Δ *mlh3* Δ = 75%, calculated from combined data of Nishant *et al.* 2008; Zanders and Alani 2009; Al-Sweel *et al.* 2017; Chakraborty *et al.* 2017). Thus, the absence of Pch2 increases the MutLγ-dependence of VDE-initiated COs at many loci, while decreasing the MutLγ-dependence of VDE-initiated COs *at HIS4* and of Spoil-initiated COs.

**Figure 3.**
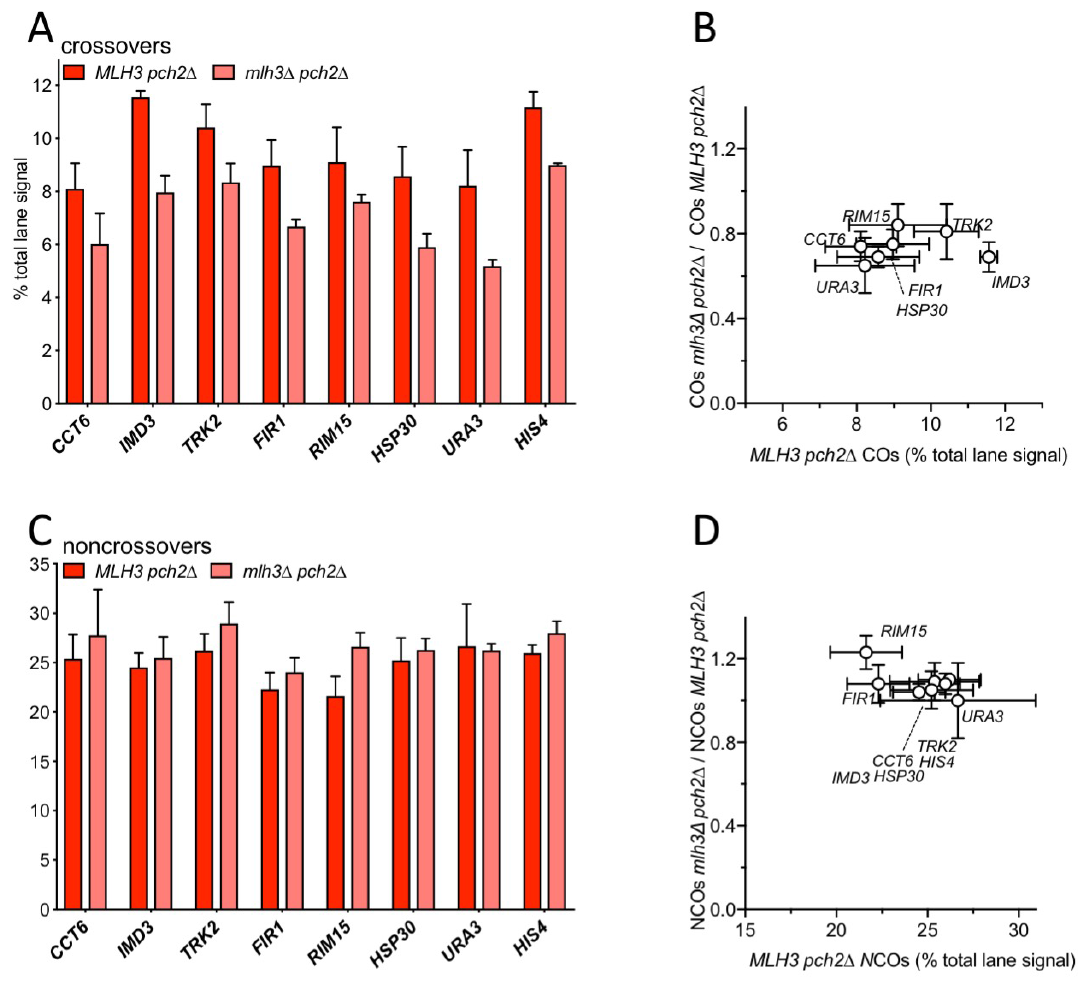
VDE-initiated crossovers in *pch2*Δ mutants are partially MutLγ-dependent. (A) VDE-initiated COs in *MLH3 pch2*Δ and *mlh3*Δ *pch2*Δ cells. (A) CO frequencies, average signal of CO1 and CO2 for 8 and 9 h samples from two independent experiments. For inserts at *CCT6, IMD3, FIR1* and *R1M15*, 9 h values are from a single experiment. Data for inserts at *HIS4* and *URA3* are from Medhi *et al.* (2016). (B) fraction of COs that are MutLγ-independent (ratio of CO frequencies in *mlh3*Δ versus *MLH3)*, plotted as a function of CO frequencies in *MLH3* strains. CO frequencies in *MLH3 pch2*Δ and *mlh3*Δ *pch2*Δ differ significantly for inserts at all loci (adjusted *p* values ≤ 0.03) except *CCT6* and *R1M15* (adjusted*p* values of 0.06 and 0.07, respectively). (C, D) VDE-initiated NCOs, as in panels (A) and (B). NCO frequencies in *MLH3 pch2*Δ and *mlh3*Δ *pch2* Δ do not differ significantly for any locus. Error bars in all panels denote standard deviation.

Spoil-initiated COs are reduced about 2-fold in mutants lacking MutLγ; this is thought to reflect unbiased JM resolution by SSNs to form both COs and NCOs, as opposed to MutLγ-mediated biased JM resolution as COs in wild type (Argueso *et al.* 2004; Zakharyevich *et al.* 2012). If the same holds true for *pch2* mutants, the ~25% reduction in COs seen in *pch2*Δ *mlh3*Δ would suggest that about half of the COs formed in *pch2 MIH3* cells are the products of MutLγ-mediated resolution, regardless of whether they were initiated by VDE or by Spoil. It therefore appears that Pch2, or factors regulated by it, prevents most VDE-initiated events from forming MutLγ-dependent COs.

### Summary and concluding remarks

In this study, we examined VDE-initiated meiotic recombination in a recombination reporter inserted at six loci in addition to the two loci (*HIS4* and *URA3*) originally examined by Medhi *et al.* (2016). With the exception of *HIS4*, VDE-initiated COs at all insert loci were largely Mlh3-independent, regardless of whether inserts were at loci in Hop1-enriched or Hop1-depleted regions of the genome. Therefore, the hypothesis proposed by Medhi *et al.* 2016, that local Hop1 occupancy determines mechanisms of JM resolution, is incorrect, at least for VDE-initiated recombination, in that it was based on analysis of inserts at a locus (*HIS4*) that appears to be exceptional.

The observation that VDE-initiated COs at most insert loci are Mlh3-independent, in turn, raises the question of whether or not VDE-initiated recombination events that occur in cells undergoing meiosis can be properly described as being “meiotic”. VDE-initiated NCOs are recovered in excess of COs (2 to 5-fold, average 3.2 ± 0.1), which is reminiscent of, although less than, the 5 to 20-fold excess of NCOs over COs seen in budding yeast mitotic recombination (Esposito 1978; Lichten and Haber 1989; Ira *et al.* 2003; Bzymek *et al.* 2010; Dayani *et al.* 2011). VDE-initiated DSB processing also resembles DSB processing in the mitotic cell cycle, in that break ends are continuously resected over time (Lee *et al.* 1998; Neale *et al.* 2002; Johnson *et al.* 2007), unlike the limited resection seen with Spo11 DSBs (Mimitou *et al.* 2017). Finally, unlike Spo11, VDE frequently cuts both sister chromatids in a single meiosis (Gimble and Thorner 1992; Gimble and Thorner 1993; Medhi *et al.* 2016), and gene CO/NCO ratio among HO endonuclease-initiated meiotic recombinants (Malkova *et al.* 2000). Further studies will be necessary to determine which of these or other factors are responsible for the marked Mlh3-independence of VDE-initiated COs at seven of the eight insert locations examined, and why the majority of VDE-initiated COs at *HIS4* are Mlh3-dependent.

In contrast, in *pch2*Δ strains, VDE-initiated COs show the same Mlh3-dependence as Spo11-initiated COs, regardless of wild-type Hop1 occupancy levels at insert loci. It therefore seems unlikely that Hop1 redistribution in *pch2*Δ mutants is the only factor responsible for the increased Mlh3-dependence of COs at most insert loci and decreased Mlh3-dependence of COs at *HIS4.* Homolog synapsis, recombinant formation and meiotic divisions are all delayed in *pch2*Δ mutants, which also display a more even distribution of the Zip1 central element protein along chromosomes and reduced CO interference (Börner *et al.* 2008; Joshi *et al.* 2009; Zanders and Alani 2009). It is possible that, in *pch2*Δ mutants, these or other defects delay the recruitment of factors necessary for MutLγ-resolution at Spoil-initiated events, and thus make them available to VDE-initiated events. Again, further studies will be necessary to test these possibilities.

In summary, the data presented here indicate that VDE-initiated recombination events are treated differently than are those initiated by Spoil during wild-type meiosis. VDE-initiated events produce an excess of NCOs over COs and, at seven of eight loci examined, form COs by MutLγ-independent mechanisms, and thus their outcome more closely resembles those of DSB repair events that occur during the mitotic cell cycle. We conclude that the full spectrum of meiotic recombination processes that occur at Spoil-initiated DSBs do not occur at VDE-initiated DSBs, and, by inference, DSBs formed during meiosis by other nucleases. Thus, our findings call for caution in the use of DSBs formed by these nucleases, or by other exogenous means, for inferring factors that control normal meiotic recombination.

## End Matter

### Author Contributions and Notes

A. S. and D. M. performed research, A. S., D. M. and M.L designed research, analyzed data and wrote the paper. The authors declare no conflict of interest.

## Acknowledgments

We thank Jean Paul Ouyan, Seyoun Kim, and Matan Cohen for help in strain construction, and Jasvinder Ahuja, Matan Cohen, Julia Cooper, and Martin Xaver for comments and discussion. This work was supported by the Intramural Research Program of the NIH through the Center for Cancer Research at the National Cancer Institute.

---

**Figure S1.**
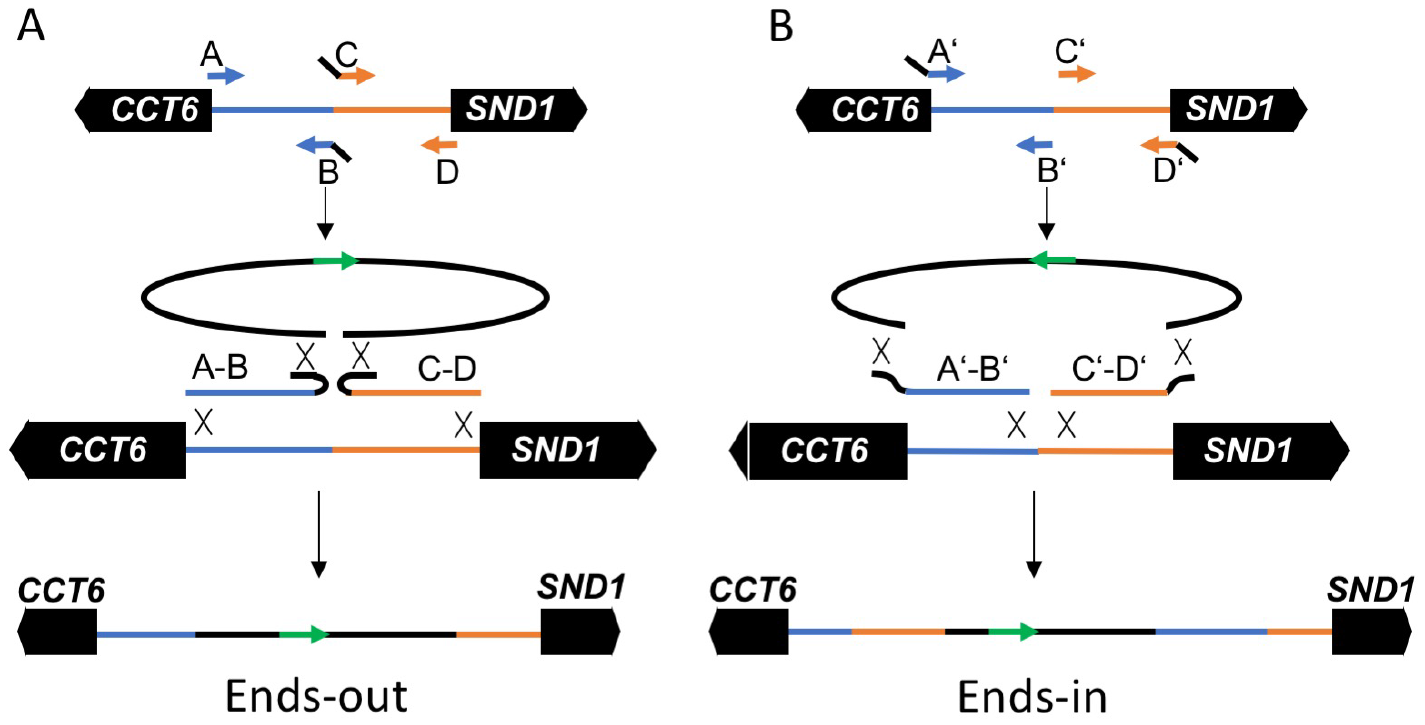
Construction of inserts. Strategy used for insertion in the *CCT6-SND1* intergenic region is illustrated. (At) Ends-out transformation, resulting in an insert that disrupts the intergenic region. PCR products corresponding to the two halves of the region are amplified, using inside primers (B and C) with homology (black lines) to the sequences to be inserted. (Ɓ) Ends-in transformation, resulting in a duplication of the intergenic region. PCR products are amplified as above, but with outside primers (A’ and D’) with homology to sequences to be inserted.

**Figure S2:**
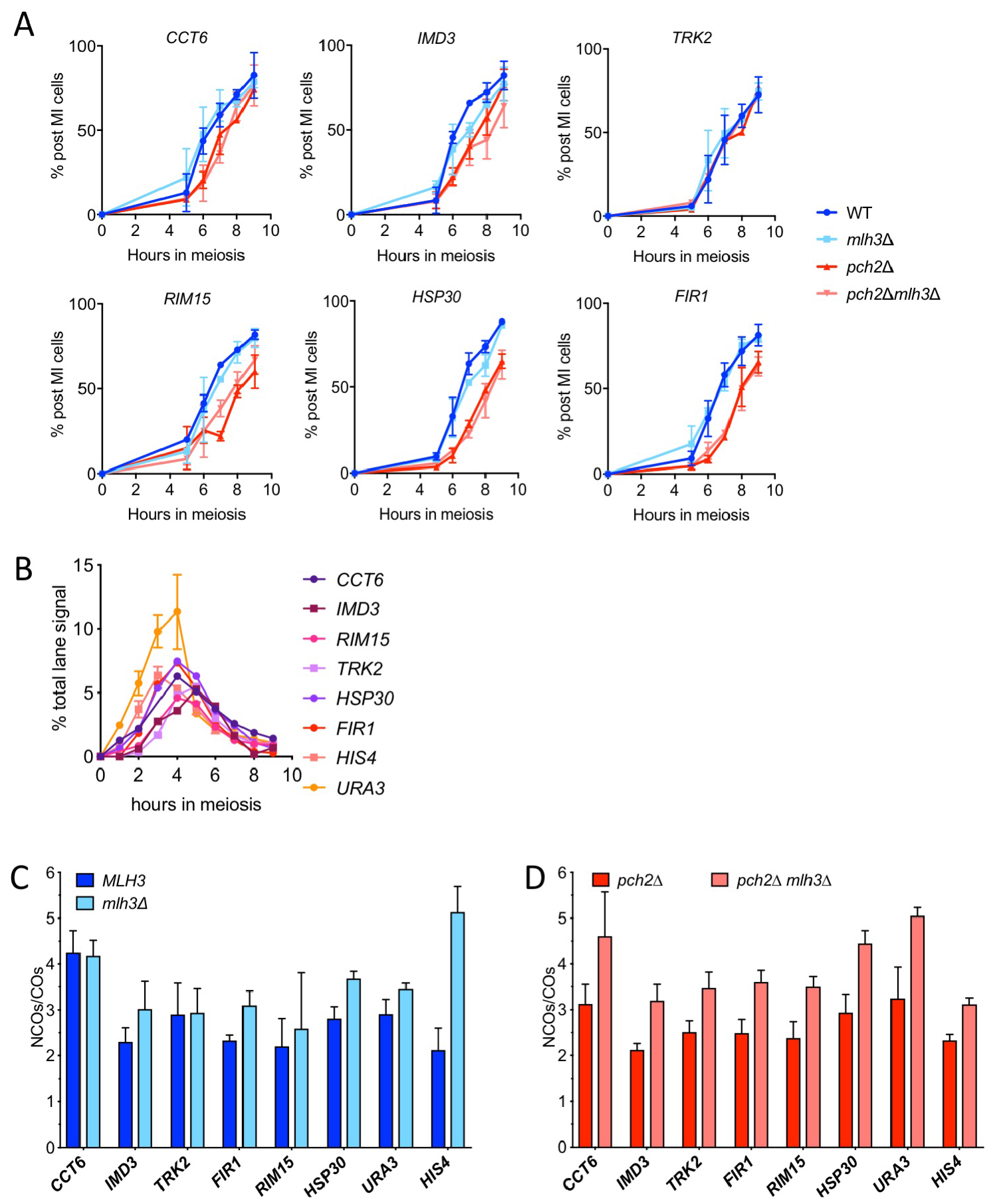
(A) Fraction of cells completing meiosis I, scored in DAPI-stained samples as cells with 2 or more nuclei. (B) VDE-initiated DSBs, scored as percent total lane signal in Southern blots containing *Ihna*dIII digests (Medhi *et al* 2016). Data for inserts at *HIS4* and *CRA3* are from Medhi et al (2016); data for all other insert loci are from a single experiment. (C) NCO/CO ratios for *PCH2* strains, calculated from mean values presented in Figure 3C, D. (D) NCO/CO ratios for *pch2* Δ strains, calculated from mean values presented in Figure 4A, B. For panels (C) and (D), eưor bars represent the sum of fractional standard deviations for each mean value. In panel (C), NCO/CO ratios in *MLH3* differ significantly from *mlh3*Δ only for inserts at *FIR1, HSP30*, and *HIS4* (adjusted *p* values of 0.02, 0.008 and 0.00001, respectively). In panel (D), NCO/CO ratios in *MLH3 pch2*Δ differ significantly from *mlh3*Δ *pch2* Δ differ significantly for all inserts (adjusted *p* values all ≤ 0.3).

**Table S1:**
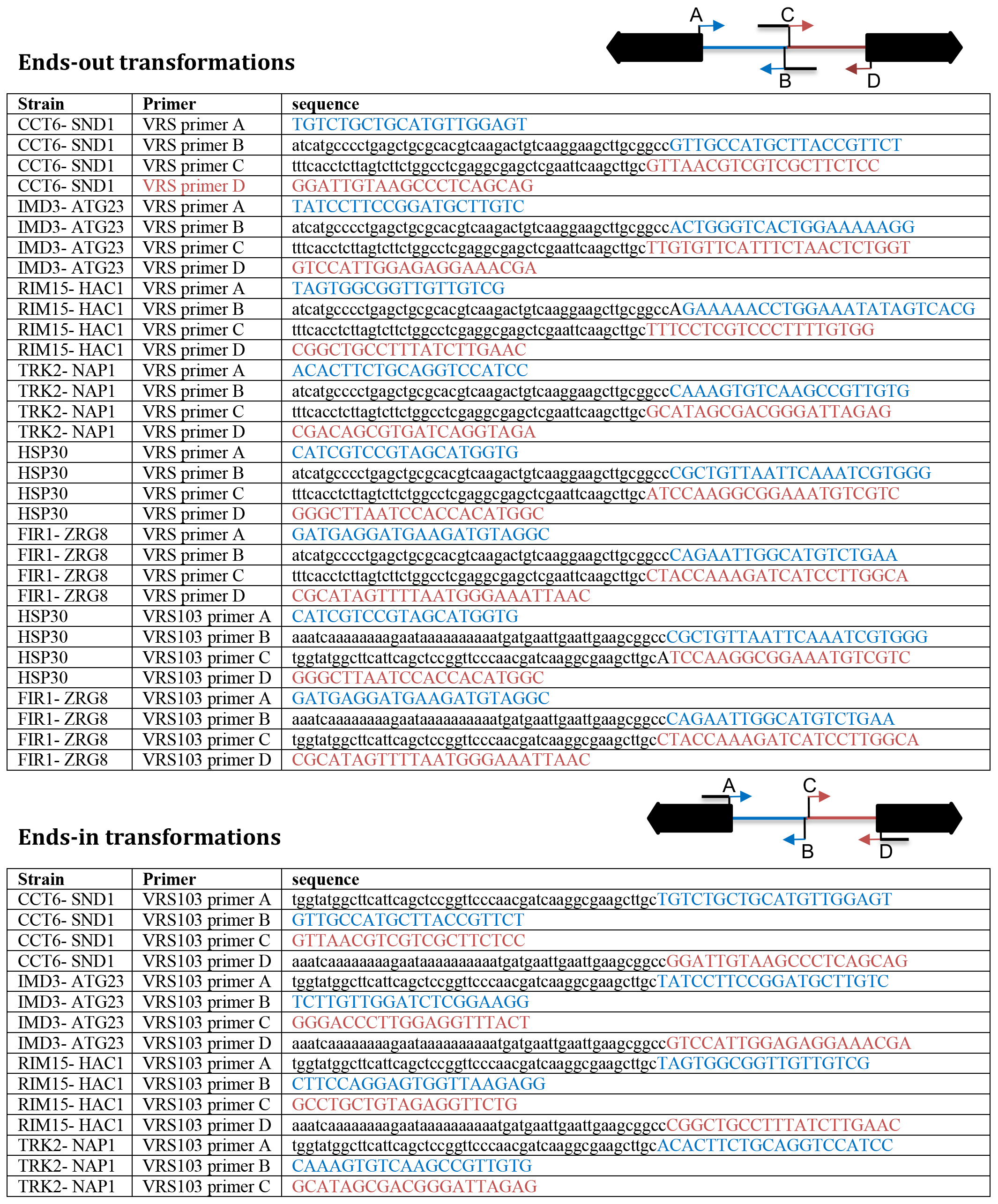
Primers for all reporter inserts. Primers are color-coded to correspond with Figure S1: black—plasmid sequences; tan or blue—corresponding yeast chromosomal sequences.

**Table S2:** Strain Genotypes. All strains are homozygous for *lys2, ho::LYS2, ura3*Δ*(hindIII-smaI), leu2, arg4*Δ*(eco47III-hpaI), trp1::hisG*, and *VMA1-103. pch2*Δ*::URA3* and *pch2*Δ*::TRP1* delete sequences between *AccI* and *BamHI* and between *AccI* and *SpeI* sites in *PCH2* coding sequences, respectively. Multiple strain names for a given genotype represent independently derived diploids, both of which were used.

| <u>Strain name</u> | <u>Relevant genotype</u> | <u>Figure</u> |
| --- | --- | --- |
| MJL3906, 3907 | *CCT6'-natMX-[arg4-VRS]-KLTRP1-'SND1* *pCUP1*<br>-----<br>*CCT6-SND1'-URA3-[arg4-VRS103]-'CCT6-SND1* *pCUP1-VDE-hygMX-pCUP1-CUP1* | 3, S1 |
| MJL3915 | MJL3907 + *mLh3Δ::KanMX/mLh3Δ::KanMX* | 3, S1 |
| MJL3944, 3945 | MJL3907 + *pch2Δ::URA3/pch2Δ::TRP1* | 4, S1 |
| MJL3952, 3953 | MJL3907 + *pch2Δ::URA3/pch2Δ::TRP1*, *mLh3Δ::KanMX/mLh3Δ::KanMX* | 4, S1 |
| MJL3908, 3909 | *IMD3'-natMX-[arg4-VRS]-KLTRP1-'AGT23* *pCUP1*<br>-----<br>*IMD3-ATG23'-URA3-[arg4-VRS103]-'IMD3-AGT23* *pCUP1-VDE-hygMX-pCUP1-CUP1* | 3, S1 |
| MJL3916, 3917 | MJL3909 + *mLh3Δ::KanMX/mLh3Δ::KanMX* | 3, S1 |
| MJL3946, 3947 | MJL3909 + *pch2Δ::URA3/pch2Δ::TRP1* | 4, S1 |
| MJL3954 | MJL3909 + *pch2Δ::URA3/pch2Δ::TRP1*, *mLh3Δ::KanMX/mLh3Δ::KanMX* | 4, S1 |
| MJL3910, 3911 | *RIM15'-natMX-[arg4-VRS]-KLTRP1-'HAC1* *pCUP1*<br>-----<br>*RIM15-HAC1'-URA3-[arg4-VRS103]-'RIM15-HAC1* *pCUP1-VDE-hygMX-pCUP1-CUP1* | 3, S1 |
| MJL3918, 3919 | MJL3911 + *mLh3Δ::KanMX/mLh3Δ::KanMX* | 3, S1 |
| MJL3948, 3949 | MJL3911 + *pch2Δ::URA3/pch2Δ::TRP1* | 4, S1 |
| MJL3955, 3956 | MJL3911 + *pch2Δ::URA3/pch2Δ::TRP1*, *mLh3Δ::KanMX/mLh3Δ::KanMX* | 4, S1 |
| MJL3912, 3913 | *TRK2'-natMX-[arg4-VRS]-KLTRP1-'FMP46* *pCUP1*<br>-----<br>*TRK2-FMP46'-URA3-[arg4-VRS103]-'TRK2-FMP46* *pCUP1-VDE-hygMX-pCUP1-CUP1* | 3, S1 |
| MJL3920, 3921 | MJL3913 + *mLh3Δ::KanMX/mLh3Δ::KanMX* | 3, S1 |
| MJL3950, 3951 | MJL3913 + *pch2Δ::URA3/pch2Δ::TRP1* | 4, S1 |
| MJL3957, 3958 | MJL3913 + *pch2Δ::URA3/pch2Δ::TRP1*, *mLh3Δ::KanMX/mLh3Δ::KanMX* | 4, S1 |
| MJL3879 | *hsp30::URA3-[arg4-VRS103]* *pCUP1*<br>-----<br>*hsp30::natMX-[arg4-VRS]-KLTRP1* *pCUP1-VDE-hygMX-pCUP1-CUP1* | 3, S1 |
| MJL3881 | MJL3879 + *mLh3Δ::KanMX/mLh3Δ::KanMX* | 3, S1 |
| MJL3976, 3977 | MJL3879 + *pch2Δ::URA3/pch2Δ::TRP1* | 4, S1 |
| MJL3970, 3978 | MJL3879 + *pch2Δ::URA3/pch2Δ::TRP1*, *mLh3Δ::KanMX/mLh3Δ::KanMX* | 4, S1 |
| MJL3880 | *FIR1'-URA3-[arg4-VRS103]-'ZRG8* *pCUP1*<br>-----<br>*FIR1'-natMX-[arg4-VRS]-KLTRP1-'ZRG8* *pCUP1-VDE-hygMX-pCUP1-CUP1* | 3, S1 |
| MJL3971, 3972 | MJL3880 + *mLh3Δ::KanMX/mLh3Δ::KanMX* | 3, S1 |
| MJL3969, 3973 | MJL3880 + *pch2Δ::URA3/pch2Δ::TRP1* | 4, S1 |
| MJL3974, 3975 | MJL3880 + *pch2Δ::URA3/pch2Δ::TRP1*, *mLh3Δ::KanMX/mLh3Δ::KanMX* | 4, S1 |

